# Chemical ecology of Himalayan eggplant variety’s antixenosis: Identification of geraniol as an oviposition deterrent against the eggplant shoot and fruit borer

**DOI:** 10.1101/2022.04.28.489959

**Authors:** Rituparna Ghosh, Dennis Metze, Maroof Shaikh, Ashish Deshpande, Dnyaneshwar M. Firake, Sagar Pandit

## Abstract

- Eggplant (*Solanum melongena*) suffers severe losses due to a multi-insecticide resistant lepidopteran pest, shoot and fruit borer (SFB, *Leucinodes orbonalis*). Heavy and combinatorial application of pesticides for SFB control renders eggplant risky for human consumption.
- We observed that 1) ovipositing SFB females can find even solitary plants of susceptible varieties and 2) they do not oviposit on Himalayan eggplant variety RC-RL-22 (RL22). We hypothesized that the olfactory cues influence ovipositing female’s host choice.
- To find these cues, leaf volatile blends of seven eggplant varieties were profiled using GCMS. Seven compounds were present in >2.5-fold concentrations in RL22 than the other varieties. In choice assays, oviposition deterrence efficacies of these candidate compounds were independently tested by their foliar application on SFB-susceptible varieties. Complementation of geraniol, which was exclusively found in RL22, reduced oviposition (>90%). To validate geraniol’s role in RL22’s SFB-deterrence, we silenced RL22’s geraniol synthase gene using virus-induced gene silencing. Geraniol biosynthesis suppression rendered RL22 SFB-susceptible; foliar geraniol application on the geraniol synthase-silenced plants restored oviposition deterrence.
- We infer that geraniol is RL22’s SFB oviposition deterrent. The use of natural compounds like geraniol, which influence the chemical ecology of oviposition can reduce the load of hazardous larvicidal pesticides.

## Introduction

Interactions of herbivore insects with their hostplants are often influenced by the plant chemistry, which drives the evolution of differential host preference and host specialization in insects. Several butterflies and moths which indiscriminately feed on the floral nectars of various species, selectively oviposit on the host species of their specialist herbivore larvae. Ovipositing females often distinguish between hosts and nonhosts with the help of these plants’ volatile organic compounds (VOCs) (Witzgall *et al*., 2005; Thöming *et al*., 2014). Generally, host VOCs act as attractants and non-host ones as repellents. Insects’ remarkably sensitive olfactory systems provide fine resolution of the complex vegetation’s odorscape during the flight (Tasin *et al*.,2005; Najar-Rodriguez *et al*., 2010; Pickett *et al*., 2012; Leppik & Frérot, 2014). They can differentiate between background host odors to find their host (Schröder & Hilker, 2008; Karlsson *et al*., 2009; Thöming *et al*., 2014). This VOC-mediated interaction is so fine-tuned in some plant-insect systems that females refrain from ovipositing on plants infested by the conspecific larvae, as these plants release repellent VOCs (De Moraes *et al*., 2001; Qiao *et al*., 2018). Furthermore, herbivory-induced VOCs of some plant species attract the natural enemies of the herbivore; this phenomenon is regarded as a natural biological control (De Moraes *et al*.,1998; De Boer *et al*., 2008; Qiao *et al*., 2018). Over the last few decades, several researchers have recommended the exploitation of these chemical ecology aspects of host selection for the management of insect pests (Pickett *et al*., 1997; Khan *et al*., 2000; Aldrich *et al*., 2003; Khan *et al*., 2008; James *et al*., 2012; Saha & Chandran, 2017; Cusumano *et al*., 2020).

Today, the protection of several crops depends on the heavy use of synthetic pesticides which pose a serious threat to human health (NRC, 1993; Bissdorf, 2010; Alavanja *et al*., 2013; Mebdoua, 2018). Pesticides also deteriorate the environment by polluting soil, water and air (Ucar & Hall, 2001; Devine & Furlong, 2007; Vymazal & Březinová, 2015; Hladik *et al*., 2018). To minimize the hazards of pesticides, chemical ecology-based eco-friendly pest management solutions are highly desired.

Eggplant (*Solanum melongena* L.; Solanaceae) is one of the crops which requires a serious and urgent reduction in pesticide application. Popularly called the king of vegetables, eggplant is the third most consumed solanaceous vegetable after potato and tomato (Chantella, 2011; Caruso *et al*., 2017; Cristol, 2018; Chapman, 2019). It is one of the nutritionist-recommended vegetables with low calorie and fat content but high antioxidant, fiber, folic acid, calcium, phosphorus, potassium and vitamins B and C content and various medicinal properties like analgesic, anti-pyretic, anti-inflammatory, anti-asthmatic and spasmogenic (Daunay, 2008; Das & Barua, 2013; Caruso *et al*., 2017; Gürbüz *et al*., 2018; Stommel & Whitaker, 2019). From the farmers’ point of view, eggplant is easy to grow, offers many varieties to suit vastly diverse geo-climatic zones that yield remarkably well in these different zones and is profitable even when cultivated on small scales (Nakasuji & Matsuzaki, 1977; FAO, 2003; Frary *et al*., 2007; Singla *et al*., 2018). Due to these virtues, the area under eggplant cultivation is rapidly increasing (Chapman, 2019). Unfortunately, eggplant is one of the heaviest pesticide-applied vegetables. It receives up to 140 sprays per ~6-month season with a frequency as high as two per day (Rashid *et al*., 2003; Del Prado-Lu, 2015; Shelton *et al*., 2019). Although eggplant is attacked by >35 pests, the eggplant fruit and shoot borer (SFB, *Leucinodes orbonalis* Guenee, Lepidoptera: Pyralidae) is mainly responsible for this high pesticide application. SFB attacks the vegetative as well as reproductive stages of eggplant and its infestation can be so severe that it can cause 45-100% loss of the marketable yield (FAO, 2003; Singla *et al*., 2018; Reshma *et al*., 2019). Most of the economic damage is inflicted by its frugivorous larval stage, which bores tunnels and deposits frass in fruits (Hautea *et al*., 2016). Since these larvae remain concealed in the fruit for most of their lifetime, several, especially non-systemic, pesticides show inadequate effects on them (FAO, 2003; Choudhary & Gaur, 2009). Due to such concealed habit, larvae are exposed to sublethal doses of pesticides which facilitates the pesticide resistance development in them (Choudhary & Gaur, 2009). Indeed, SFB has developed resistance against commonly used pesticides of several different classes in a short time (Table S1). Consequently, various synthetic pesticides are often used in heavy doses and combinations for its control (Choudhary & Gaur, 2009; Rahman, 2009; Reshma *et al*., 2019). Such pesticide abuse endangers the health of farmworkers, pollinators and other non-target insects in the eggplant agroecosystem and can also cause a resurgence of secondary pests. Moreover, residues of these pesticides render eggplant harmful for human consumption (Kalawate & Dethe, 2011; Dasika *et al*., 2012; Del Prado-Lu, 2015; Shelton *et al*., 2019).

Assorted means of attaining SFB control have been explored to lessen the dependence on pesticides; for example, screening and breeding for resistant or tolerant varieties (Lal, 1991; FAO, 2003; Prabhu *et al*., 2008; Khan & Singh, 2014; Kassi *et al*., 2019), introgression of resistance traits from wild relatives (Dhankhar *et al*., 1982; Dhankar, 1988; Kumar & Sadashiva, 1996; Rotino, 1997), finding biophysical and biochemical bases of resistance traits for their introduction in susceptible varieties (Naqvi *et al*., 2009; Khorsheduzzaman *et al*.,2010; Wagh *et al*., 2012; Prasad *et al*., 2014; Nirmala & Vethamoni, 2016), intercropping with deterrent herbs (Khorsheduzzaman *et al*., 2010; Calumpang *et al*., 2013) and application of botanicals (Adiroubane & Raghuraman, 2008; Calumpang & Ohsawa, 2015), sex pheromones (Zhu *et al*.,1987; Cork *et al*., 2005), natural enemies (Naresh *et al*., 1986; Sasikala *et al*., 1999; FAO, 2003) and entomopathogens (Beevi & Jacob, 1982; Khorsheduzzaman *et al*., 1998; Mainali *et al*.,2013). Outcomes of most of these were used in the integrated pest management; however, none of them could appreciably control SFB (Alam *et al*., 2003; Mainali, 2014; Lalita & Kashyap, 2020). Consequently, the transgenic eggplant was created to control SFB by incorporation of a gene coding for the insecticidal Cry1Ac protein from *Bacillus thuringiensis* (*Bt*) (Choudhary & Gaur, 2009; Kumar *et al*., 2011; Hautea *et al*., 2016). Although these transgenics showed the desired resistance against SFB, acceptance to them remained limited due to socioethical concerns (Krishna & Qaim, 2007; Seetharam, 2010; San-Epifanio, 2017; Glaab & Partzsch, 2018).

Here we present a study in which, two observations on SFB’s host selection behavior led to a molecular and chemical ecology investigation of host-SFB interaction revealing the chemistry of eggplant’s resistance against this tenacious pest. While collecting SFB insects from eggplant fields for initiating a laboratory culture, we found eggs and larvae even on solitary plants that were far away from eggplant fields. Secondly, we observed that one of our germplasm collections from the Eastern Himalayas showed extremely low or no SFB oviposition. Together, these observations suggested that ovipositing females’ host finding and selection abilities were key to SFB’s occurrence. We hypothesized that eggplant’s VOC blend characteristics influence SFB’s host selection. We tried to find the putative deterrents from the Himalayan variety. Lastly, we validated our findings by the reverse genetics-based characterization of the repellent biosynthesis gene in this variety.

## Materials and methods

### Eggplant field

Eastern Himalayan variety RC-RL-22 (RL22) [IC-0634845; National Bureau of Plant Genetic Resources (ARIS Cell), India] and six popular Indian eggplant varieties Ankur Kavach (KV), Ankur Vijay (VJ), JK 6829 (JK), Riccia Hirvi kateri (HK), KGN’s pinstripe (KP) and Omaxe CVK MK 124 (CVK) were used in this work (Fig. 1a). All seven varieties were planted in the experimental field of Indian Institute of Science Education and Research (IISER), Pune (18.547669 ° N, 73.807636 ° E) in a randomized complete block design (Fig. 1b) with four blocks and each one containing 5 individuals of the abovementioned plants (n= 20 plants per variety, in four blocks) with 1 m spacing between individuals. Manures and fertilizers were provided as recommended for this region (Anonymous, 2010). No pesticides were applied in the field.

**Fig. 1.**
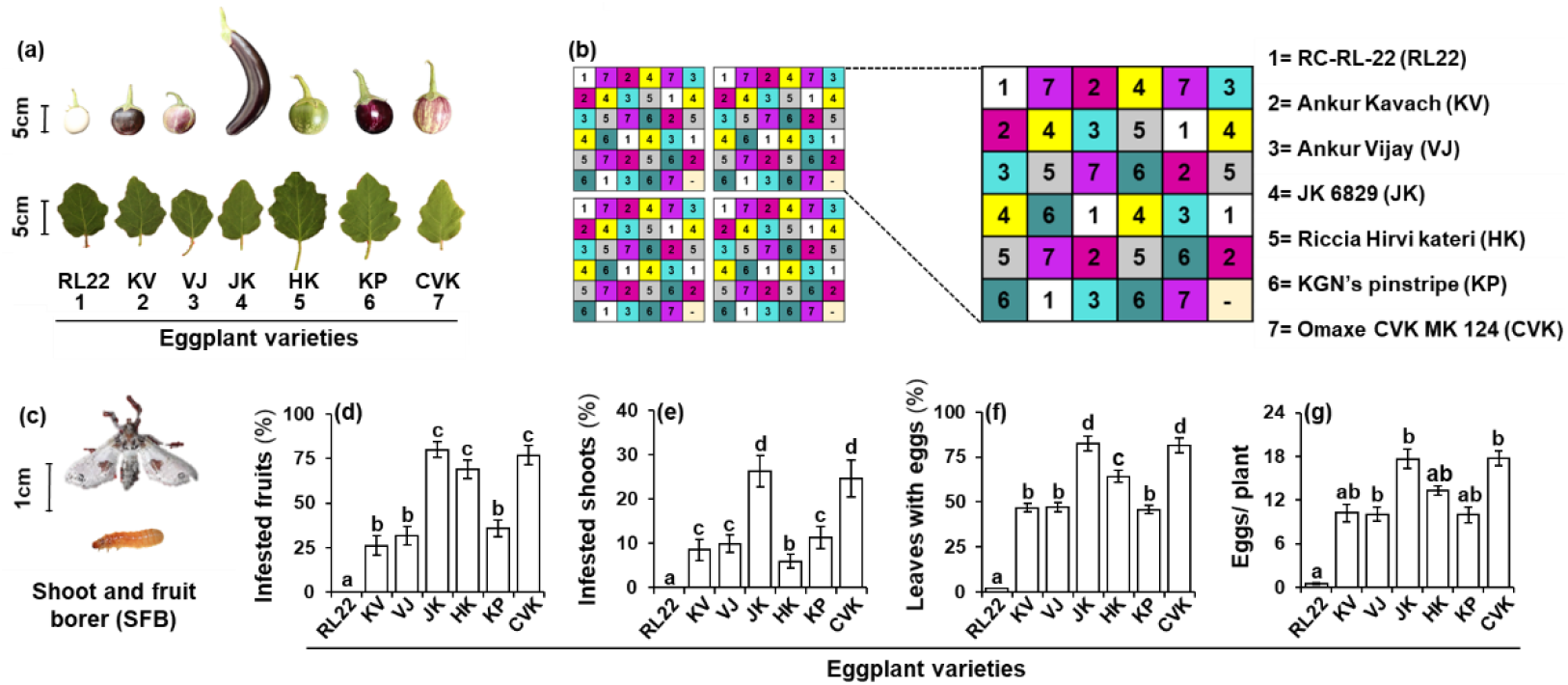
SFB does not prefer Himalayan eggplant variety RL22. (a) Fruits and leaves of seven eggplant (*Solanum melongena*) varieties, eastern Himalayan eggplant variety RL22 and six others (KV, VJ, JK, HK, KP and CVK) used in this study. (b) Schematic of experimental field with the randomized complete block design. (c) Moth and larva of SFB (*Leucinodes orbonalis*). SFB infestation (%) in (d) fruits (*F*= 133.7, *df*= 50.67, *P*< 0.0001) and (e) shoots (*F*= 25.65, *df*= 50.67, *P*< 0.0001) of seven varieties, indicating that RL22 is the least preferred. (f) Oviposition preference based on leaves (%) harboring eggs (*F*= 280.5, *df*= 54.89, *P*< 0.0001) and (g) number of eggs laid per plant (*F*= 134.6, *df*= 53.49, *P*< 0.0001) on different varieties. Significant differences (*P*≤ 0.05) were determined using Welch F test with Games-Howell *post hoc* test.

### Insects

SFB (Fig. 1c) eggs, larvae and pupae were collected from the eggplant fields in and around Pune to initiate the laboratory culture. Larvae were maintained in aerated polypropylene containers [(l) 30 cm× (b) 20 cm× (h) 10 cm] incubated inside a climate chamber (26± 1°C temperature, 65± 5% relative humidity, 16 h light and 8 h dark photoperiod) and were reared on artificial diet (Salunke *et al*.,2014). Pupae were maintained in the dark. For mating, moth pairs were kept in jars [(id) 10 cm× (h) 20 cm] containing healthy eggplant twigs as the oviposition substrates and were fed 10% (w/v) aqueous sucrose solution with the help of a cotton wick. Adults from the third generation of this culture were used in the experiments.

### Natural occurrence of SFB in the field

The incidence of SFB on seven eggplant varieties in the randomized complete block design was measured. On each plant, the number of SFB-infested fruits and shoots were counted. To understand whether the ovipositing SFB females showed differential preferences to the seven eggplant varieties, percentage of leaves (per plant) harboring eggs and number of eggs per plant were recorded for all the varieties.

### Determining the factors influencing the ovipositing females’ host selection

#### Multiple-choice assays

All the insect behavioral assays were performed inside the climate chamber using potted (800 cm^3^ pots) juvenile plants with five fully expanded leaves. In a multiple-choice oviposition assay, one plant of each eggplant variety was used. Besides, one artificial plant (AP) having five leaves made of a Whatman filter paper (grade 237 equivalent to grade 3 CHR; Sigma-Aldrich, India) was also used as an inert control. These plants were arranged inside a nylon mesh (160 μm) tent with a 30 cm interplant distance. One SFB female, which had mated 18-24 h before the assay, was released in this tent and allowed to oviposit for 10 h. A cotton wick dipped in a sucrose solution was provided to moths for feeding during the assay. The assay was repeated 20 times with a different randomized hostplant arrangement every time to negate any hostplant positional effects. At the end of each assay, eggs on each plant were counted.

#### Multiple-choice assays using VOC blend-complemented APs

Since it is well known that SFB females routinely oviposit on eggplant leaves, irrespective of the presence of flowers and fruits on plants (Prabhat & Johnsen, 2000; Ardez *et al*., 2008; Mannan *et al*., 2015), we hypothesized that the putative attractants or repellents were mainly from leaves. To find whether the putative attractants from the SFB-preferred hosts and putative repellants from the SFB-disliked hosts were present in their VOC blends, we conducted multiple-choice assays using APs and complemented them with the VOC blend extracts of different varieties. The cumulative area of abaxial and adaxial surfaces of each filter paper leaf was 100± 10 cm^2^ and its weight was 1.0± 0.1 g, which was similar to the average surface area and the average weight of eggplant leaves. For complementation, VOC blends of the seven varieties were obtained as follows. 3.0± 0.5 g of freshly harvested fully expanded healthy leaves were immersed in 20 ml of GCMS grade dichloromethane (DCM; RANKEM, India) and extracted for 6 hours by rolling on a tube roller. Leaves were discarded and the DCM extract was dehydrated using anhydrous sodium sulfate (Sigma-Aldrich, India). The leaf VOC extract of each eggplant variety was coated on the leaves of different APs in such a way that the extract from 1g eggplant leaf was used for every 1g of filter paper leaf. DCM was allowed to evaporate for 10 min and then females were released in the assay set-up. DCM-complemented plants were used as controls. Assays were conducted and results were analyzed using the same procedure as assays using living plants, mentioned above.

#### Dual-choice assays to determine the effects of individual VOCs

To determine whether the candidate VOCs (those found either only in RL22 or significantly higher in RL22 than other varieties) repelled SFB females, a series of dual-choice assays were conducted. These assays were conducted in two ways: 1) using APs and 2) using plants of all seven varieties. In the plant-based assay, two individuals of a variety were taken; one was complemented with the candidate compound (100μl g^-1^; diluted in its solvent) and the other with the candidate compound’s solvent (100μl g^-1^). These control and test plants were kept in a nylon mesh tent inside the climate chamber into which, one mated female was released and allowed to oviposit; these assays were conducted for 6 h. Plants of six eggplant varieties were complemented to match the candidate compound’s physiological concentration (nmol g^-1^) in RL22 (mean nmol g^-1^). After the assay, the eggs were counted. For each candidate compound, twenty such assays were conducted with each variety. Pure standards of these compounds used in complementation assays were obtained from Sigma-Aldrich, India. Compounds o-tolylcarbinol and p-tolylcarbinol could not be procured and therefore, were not used for assays.

#### No-choice assays to determine repellant’s effective concentration

After determining the repellent compound, to determine its optimally effective concentration, serially increasing concentrations were coated on leaves of the highly preferred CVK variety and AP. By considering the physiological concentration of the repellent compound in RL22 (nmol g^-1^) as 1X, CVK and APs were complemented to attain 0.5, 1.0, 1.5, 2.0 and 2.5X concentrations. These plants were presented to the mated females for oviposition and the number of eggs laid on each choice was enumerated.

#### Dual-choice assays using *Sm*GS-silenced plants

Dual-choice assays were performed using mated females as described above. Females were provided a choice between 1) *Sm*GS-silenced and WT, 2) *Sm*GS-silenced and empty pTRV2 vector transformed (EV), 3) *Sm*GS-silenced and *Sm*GS-silenced geraniol-complemented plants.

### Characterization and quantification of leaf VOCs using gas chromatography (GC)

For GC-based leaf VOC analysis, fully expanded leaves were collected (n= 6 plants per variety or treatment) between 9.00 pm and 1.00 am, which is the peak oviposition time of SFB females (Mannan *et al*., 2015). VOCs were extracted from the leaves using our previously described solvent extraction method (Pandit *et al*., 2009). Qualitative and quantitative analysis of the extracted volatiles was conducted using 7890B GC and 7000D triple quadrupole mass spectrometer (MS) system coupled with a flame ionization detector (FID) (Agilent Technologies, India). GCMS system was used for the identification of hostplant VOCs by comparing their mass spectra with the ones in mass spectral databases (Integrated Wiley Registry 11th Edition-NIST 2017 Mass Spectral Library) using the NIST mass spectral search program (ver. 2.0). Sample extract (2 μl) was injected in a spitless injection mode via multimode autosampler and compounds were separated on a DB5 column (30 m l× 0.32 mm ID× 0.25 μm film thickness) (Agilent J&W Scientific, India). Inlet temperature was constant at 250 °C and carrier helium gas flow was 2 ml min^-1^. The GC oven program was as follows: the temperature was held for 1.5 min at 40 °C, increased to 180 °C at the rate of 2.5 °C min^-1^, increased to 280 °C at the rate of 20 °C min^-1^ and held at 280 °C for 5 min. MS parameters were: electron impact (EI) ionization with 70 eV energy and scanning m/z 30-600 at scan speed 7 cycle sec^-1^. For GCFID, which was used for the quantification of volatiles, GC parameters were the same as above. Detector temperature was maintained at 250 °C. Relative quantification of all the VOCs was done by normalizing the peak area of the analytes with that of the internal standard nonyl acetate (Sigma-Aldrich, India).

### Isolation of geraniol synthase gene (*Sm*GS) from RL22

To identify the monoterpene synthase gene responsible for geraniol production in *S. melongena*, we first obtained known GS sequences of Solanaceae members from the NCBI database. *Petunia* x *hybrida* (MK159028.1), the only Solanaceae GS with complete cds, was used as a query to search Eggplant Genome Project Database (Barchi *et al*., 2019) and Sol Genomics Network (Fernandez-Pozo *et al*., 2015) using Blastn. A complete cds of SMEL_001g121430.1.01, which showed 74% identity with the *Petunia* x *hybrida* GS was used as a candidate (*Sm*GS) for the further reverse genetics characterization. Other eggplant monoterpene synthases *Sm*MTPS1 (Sme2.5_12717.1_g00002.1) and *Sm*MTPS2 (SMEL_001g121460.1.01), which were highly similar (92% and 76%, respectively) to the candidate geraniol synthase *Sm*GS were used to test whether the *Sm*GS silencing was specific and did not cause any off-target co-silencing.

### Virus-induced gene silencing (VIGS) of *Sm*GS

To ascertain that RL22’s geraniol was responsible for the repellence of ovipositing SFB females, we conducted a reverse genetics-based functional characterization of the *Sm*GS using VIGS. Gene silencing strategy should be designed in such a way that siRNA must not co-silence the non-target genes. For this, no ≥20b stretch of the candidate mRNA section selected for siRNA generation should be 100% similar to the homologous regions of the non-target mRNA (Kumar *et al*., 2012). *Sm*GS ORF section not showing such ≥20 b identity with the *Sm*MTPS1 and *Sm*MTPS2 transcripts could not be found. Therefore, along with a 91 bp of the 3’-section of *Sm*GS ORF, we used a (153 bp) section of *Sm*GS 3’-UTR. This 244-bp *Sm*GS fragment was amplified from the leaf cDNA (primers given in Table S2). It was cloned in the pTRV2 vector (Liu *et al*., 2002; Liu *et al*., 2012) in an antisense orientation to generate the pTRV2-*Sm*GS construct. pTRV1, pTRV2 and pTRV2-*Sm*GS constructs transformed in *Agrobacterium tumefaciens* strain GV3101 were independently infiltrated into eggplant leaves, as described by Saedlar and Baldwin (2004). After four weeks of infiltration, plants were used for choice assays and leaves were collected for analyzing the target *Sm*GS transcript abundance and geraniol concentrations. In all the VIGS experiments, empty pTRV2 vector (EV) transformed plants and wild type RL22 (WT) plants were used as negative controls (Kumar *et al*., 2012).

### RNA isolation and quantitative real-time PCR (qPCR)

VIGS-mediated silencing of *Sm*GS gene was confirmed using qPCR. Leaves harvested from pTRV2-*Sm*GS, EV, and WT plants were flash-frozen in liquid nitrogen and were stored at −80 °C until further use. RNA was isolated from 500 mg pulverized tissue using RNAiso Plus reagent (Takara, Japan), as recommended by the manufacturer. cDNA was synthesized using the PrimeScript Reverse Transcriptase kit (Takara, Japan) as per the manufacturer’s instructions. To determine *Sm*GS silencing efficiency, qPCRs were performed using SYBR Premix Ex Taq II reagent kit (Takara, Japan) and CFX96 Touch Real-Time PCR Detection System (Biorad, USA). Thermocycling conditions were: 95°C for one minute, 39 cycles of 95°C for 45 seconds, 60°C for 45 seconds, 72°C for 45 seconds. The presence and abundance of virus were analyzed with the help of qPCR of viral coat protein transcripts. To determine whether the closely related gene was co-silenced, transcripts of *Sm*MTPS1 and *Sm*MTPS2 were analyzed. For calculating the relative abundance of these transcripts, Cyclophilin A, a housekeeping gene coding for a peptidyl-prolyl isomerase was used as an internal reference (Kanakachari *et al*., 2016). Details of all the primers used in these analyses are given in Table S2. All results were obtained from six biological and two technical replicates.

### Statistical analyses

In the field experiments involving the randomized complete block design, data from each block were analyzed cumulatively as well as separately. The homogeneity of quantitative data (mean± SE) was tested using Levene’s test. Homogenous data were analyzed by one-way ANOVA and the statistical significance was determined by Tukey’s *post hoc* test (*P*≤ 0.05). Normal non-homogenous data were analyzed using Welch ANOVA and Games-Howell *post hoc* test (*P*≤ 0.05). Non-parametric data were analyzed using Kruskal-Wallis test and Dunn’s *post hoc* test with Bonferroni correction (*P*≤ 0.05). Data of dual-choice assays were analyzed using Student’s t-test (2-tailed, *P*≤ 0.05). For this, quantities of compounds not detected in one or more varieties (mentioned as ND in Table S3) were considered to be zero.

We used correlation analysis to understand whether a female’s preference of eggplant variety for oviposition influences the larval occurrence in shoots and fruits of different varieties. Directions and strengths of correlations between the natural abundance profiles of larvae in shoots and fruits, eggs and host preferences of ovipositing females in multiple-choice assays were calculated using Spearman’s Rho (*r_s_*) tests. The significance of obtained correlations was determined using a 2-tailed test (*P*≤ 0.05).

Principal component analysis (PCA) was performed to understand the relation of different eggplant varieties based on their VOC composition using multivariate statistical package 3.1 (MVSP, Kovach Computing Services, 2020). PCA was performed on the quantitative VOC data in standardized mode using a correlation matrix and by applying Kaiser’s rule. To understand the influence of various attributes of the character (quantity or uniqueness) on the ordination, PCA was also performed with the transposed data.

## Results

### Himalayan eggplant variety RL22 shows very low SFB oviposition and infestation in the field

To assess whether SFB larvae show differential occurrence on different varieties, fruits and shoots of all plants in our field were surveyed. JK (80.07± 4.33%), CVK (76.85±5.32%), and HK (68.98± 5.03%) showed the highest fruit infestation, followed by KP (35.66± 4.75%), VJ (31.56± 5.15%) and KV (26.12± 5.39%) (Fig. 1d). RL22 showed no SFB infestation in the 472 examined fruits (Fig. 1d). Shoot boring (Fig. 1e) was highest in CVK (24.71± 4.23%) and JK (26.27± 3.55%), followed by KP (11.41± 2.48%), VJ (9.94± 1.94%), KV (8.53± 2.36%) and HK (5.96± 1.55%). No shoot infestation was found in the 988 inspected shoots of RL22 (Fig. 1e).

We also tried to understand 1) whether the ovipositing SFB females’ resolution of the field is robust even when it contains mixed cultivation of multiple eggplant varieties, 2) whether they can differentiate between different eggplant varieties and 3) whether they can identify RL22, which was found to be the least preferred variety in the preliminary observations, even in a mixture of eggplant varieties. For this, we analyzed the oviposition in the randomized plantation. When we tried to analyze the females’ host preferences based on the percentage of leaves of each variety they selected for oviposition, a similar trend was observed; CVK (81.47± 4.07%) and JK (82.62± 4.13%) were found to be the most preferred eggplants for oviposition, followed by HK (64.25± 2.78%), VJ (47.13± 2.35%), KV (46.75± 2.33%), and KP (45.8± 2.29%) (Fig. 1f). Congruent with the preliminary observation from its native habitat, preference to RL22 was found to be the least (2.28± 0.11%) preferred host. Most numbers of eggs (per plant) were found on CVK (17.75± 1.04) and JK (17.65± 1.33) followed by HK (13.35± 0.65), KV (10.2± 1.22), VJ (10.05± 0.94), and KP (9.95± 1.02) (Fig. 1g). Very little (0.5± 0.18) oviposition was observed on RL22 (Fig. 1g). Same SFB preference trends were observed when these data from different blocks were separately analyzed (Fig. S1a-p).

Since the occurrence of eggs showed a similar trend to that of larval occurrence, we attempted to find whether the larval occurrence was correlated to that of eggs. Shoot and fruit infestation of SFB larvae on seven eggplant varieties, which showed strong positive correlation with each other (*rs*= 0.79, *P*= 0.04) also showed strong positive correlations with the oviposition profile (*rs*= 0.86, *P*= 0.01 and *rs*= 0.64, *P*= 0.1, respectively). Such strong positive correlations indicated that the differential larval occurrence across eggplant varieties could be attributed to the differential oviposition.

### Even in the controlled environment, SFB females do not oviposit on RL22 or APs complemented with its VOC blend

To ascertain that the field-observed preference of SFB was not influenced by the field’s environmental factors, we conducted multiple host choice assays with the ovipositing females in a controlled environment (Fig. 2a). Similar to the field results, when mated females were presented with a choice of the seven eggplant varieties, the highest oviposition was observed on CVK (21.35± 2.73) and JK (18.8± 2.12), followed by VJ (7.75± 1.96), KP (7.1± 2.33), KV (6.95± 1.22) and HK (6.84± 1.75) (Fig. 2b). Females laid a few eggs on AP (0.65± 0.27) but laid no eggs on RL22 (Fig. 2b).

**Fig. 2.**
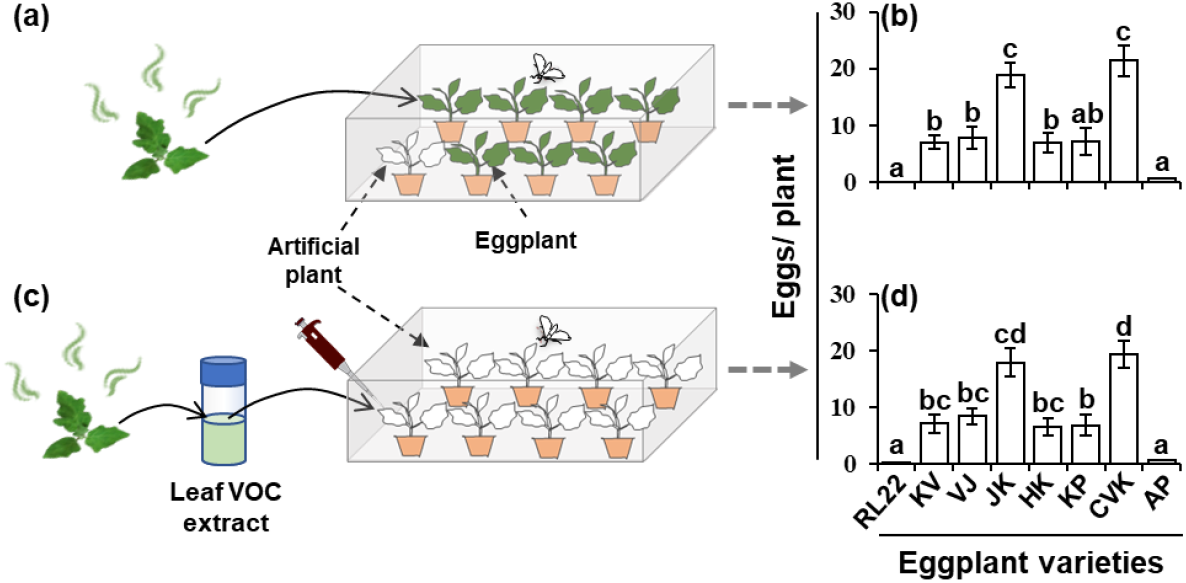
Ovipositing SFB’s host preference observed in the field remains unchanged in the controlled environment. (a) A schematic of multiple-choice assays conducted to analyze ovipositing SFB (*Leucinodes orbonalis*) females’ host preference, using eggplants and a control artificial plant (AP; Fig. S2a). (b) Eggs (mean± SE) laid on different plants used in the assay (*F*= 29.16, *df*= 57, *P*<0.0001). (c) A schematic of multiple-choice assays conducted using APs complemented with the VOC extracts of different eggplant (*Solanum melongena*) varieties. (d) Eggs (mean± SE) laid on different VOC extract-complemented APs used in the assay (*X*^2^= 88.09, *P*< 0.0001). Significant differences (*P*≤ 0.05) in (c) were determined using Welch F test with Games-Howell *post hoc* test and in (d) Kruskal-Wallis test followed by Dunn’s *post hoc* test with Bonferroni correction.

We wanted to determine if RL22’s repellence could be attributed to its VOCs or other characters such as morphological features. For this VOC blend extracts of different varieties were separately coated on APs (Fig. S2a) and presented simultaneously to mated females (Fig. 2c). The highest number of eggs was found on plants complemented with the blends of CVK (19.25± 2.36) and JK (17.85± 2.44), followed by VJ (8.4± 1.46), KV (7.1± 1.59), KP (6.8± 1.86), HK (6.45± 1.51) (Fig. 2d). RL 22 (0.05± 0.05) and DCM-complemented artificial plant controls (0.6± 0.21) showed extremely low oviposition (Fig. 2d). Females did not differentiate between untreated and DCM-complemented APs (Fig. S2b).

That the ovipositing females’ host choices in the climate chamber were similar to that in the field was ascertained by the strong positive correlation between them (*rs*= 0.75, *P*<0.06; Fig. S2c). Correlation analysis also ascertained that the oviposition on VOC extract complemented APs was highly similar to that on actual plants used in the multiple-choice assays (*rs*= 1.0, *P*< 0.0004; Fig. S2c) and also to that observed on the plants in the field (*rs*= 0.75, *P*< 0.06; Fig. S2c). The similarity in oviposition preference in all these assays suggested that ovipositing SFB female’s host preference is indeed influenced by leaf VOCs; more importantly, the preference-influencing VOC was a constituent of the extract.

### RL22 leaf VOC blend is different from that of other eggplant varieties

To understand the role of leaf volatile cues in SFB female’s host selection, a detailed GCMS/ FID-based volatile analysis was conducted. A total of 21 compounds were detected (Fig. 3a, Table. S3). Total eggplant volatile content varied from 68 nmol g^-1^ in KP to 146 nmol g^-1^ in RL22. The total volatile content of RL22 was found to be significantly higher than that of all other eggplant varieties (Fig. S3a).

**Fig. 3.**
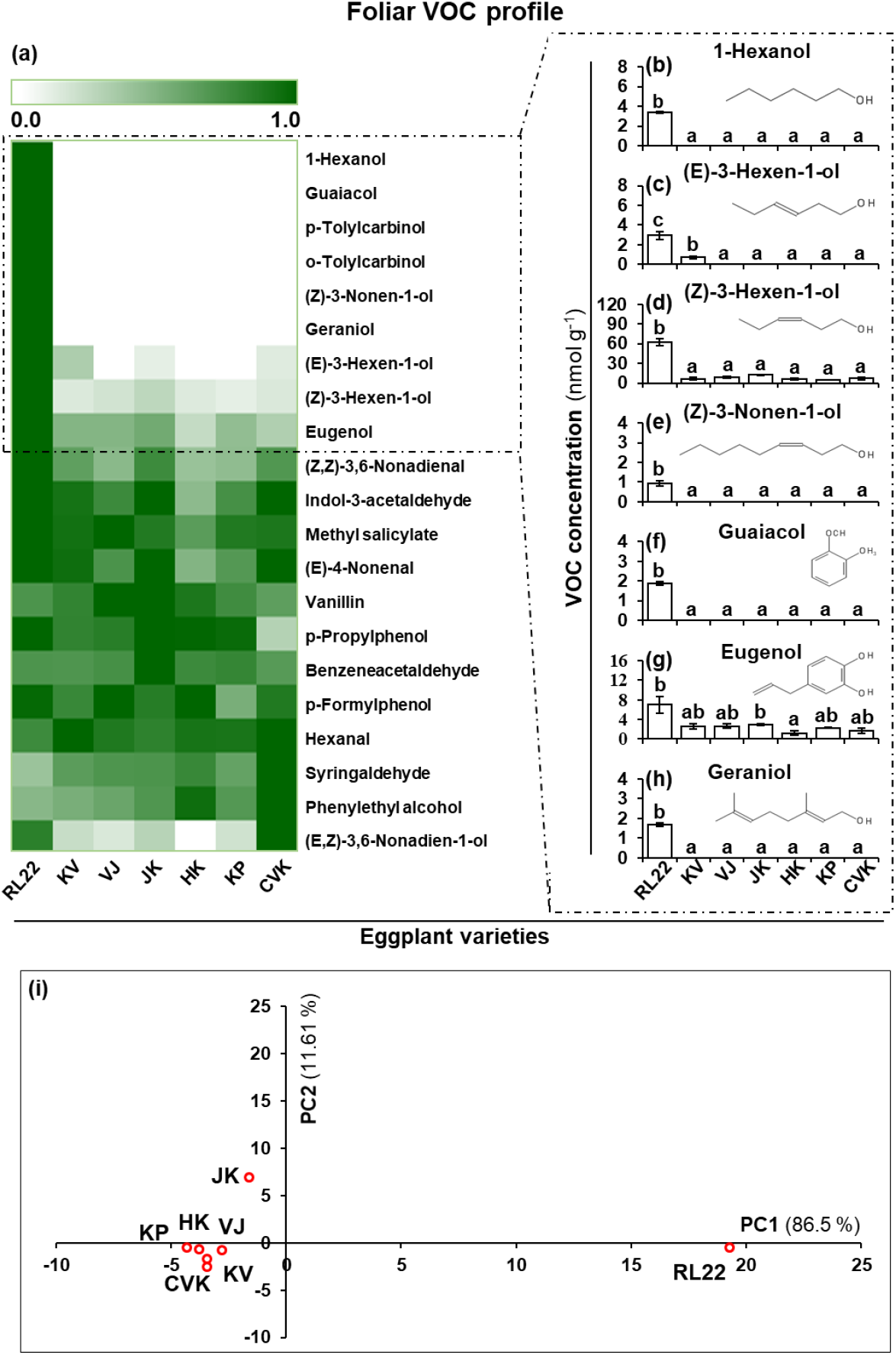
RL22’s leaf VOC profile is different from the other eggplant varieties. (a) Heatmap showing a comparison between VOC profiles of seven eggplant (*Solanum melongena*) varieties. VOCs, which were found either only in or significantly higher in RL22 blend than other varieties: (b) 1-hexanol (*F*= 58.22, *df*= 15.48, *P*< 0.0001), (c) (E)-3-hexen-1-ol (*F*= 16.06, *df*= 15.38, *P*< 0.0001), (d) (Z)-3-hexen-1-ol (*F*= 13.64, *df*= 14.05, *P*< 0.0001), (e) (Z)-3-nonen-1-ol (*F*= 4.225, *df*= 15.48, *P*= 0.01), (f) guaiacol (*F*= 7.495, *df=* 15.48, *p*= 0.0007), (g) eugenol (*F*= 5.107, *df*= 15.31, *P*= 0.005), and (h) geraniol (*F*= 7.5285, *df=* 15.48, *P*= 0.0007). Significant differences (*P*≤ 0.05) were determined using Welch F test with Games-Howell *post hoc* test. (i) A principal component analysis (PCA) score plot showing the hostplant grouping based on their VOC compositions; RL22 clearly separated from the other six eggplant varieties.

Eggplant VOC blends were found to be mainly comprised of benzene derivatives and C6 and C9 alcohols and aldehydes. Qualitatively, VOC blends of all the eggplant varieties except RL22 were quite similar. They all contained 14 compounds, which showed quantitative differences between different varieties. Henceforth, we refer to this blend as ‘common eggplant blend’ (CEB). In addition to CEB, the RL22 blend contained six compounds *viz*. 1-hexanol, guaiacol, o-tolylcarbinol, p-tolylcarbinol, (Z)-3-nonen-1-ol and geraniol, which were not detected in the other varieties; RL22 was also found to contain (Z)-3-hexen-1-ol, (E)-3-hexen-1-ol and eugenol in >5.0- and >2.5-fold higher concentrations than the other varieties, respectively (Fig. 3b-h, Table. S3). Hereafter, we refer to these nine VOCs as ‘RL22 compounds’.

The CEB was dominated by benzene derivative compounds, which constituted >60%of the total volatile content (Fig. S3b). RL22 blend contained 39.5% of benzene derivative compounds (58.16 nmol g^-1^). Major compounds were benzeneacetaldehyde, methyl salicylate, eugenol, and vanillin, of which benzeneacetaldehyde dominated the CEB (highest in JK= 43.47± 3.18 nmol g^-1^ and lowest in CVK= 22± 1.52 nmol g^-1^); its alcohol derivative phenyl-ethyl alcohol was relatively less abundant (Table S3). Eugenol concentration was significantly higher in RL22 (6.97± 1.69 nmol g^-1^) than in other eggplant varieties (Table. S3). Guaiacol was uniquely present in RL22 (1.89± 0.37) (Table. S3).

The eggplant C6 compound blend consisted of aldehyde hexanal and alcohols *viz*. (E)-3-hexen-1-ol, (Z)-3-hexen-1-ol and 1-hexanol (Fig. S3). The total C6 compound content of eggplants showed a large variation ranging from 14 nmol g^-1^ in KP and 76 nmol g^-1^ in RL22 (Fig. S3c). C6 compounds represented 34% of the RL22’s VOC blend. RL22 showed the highest abundance of (Z)-3-hexen-1-ol (61.82± 5.53 nmol g^-1^), which was >4-fold higher than that in any other eggplant variety (Fig. 3d, Table S3). C9 alcohols and aldehydes detected in the analysis were (E)-4-nonenal, (Z, Z)-3,6-nonadienal, (Z)-3-nonen-1-ol and (E, Z)-3,6-nonadien-1-ol. RL22 showed the highest C9 compound content (11.39 nmol g^-1^) among all the varieties (Fig.S3d, Table S3). Geraniol, a monoterpene alcohol, the only terpenoid detected in the eggplant blend, was exclusively found in RL22 (1.68± 0.11 nmol g^-1^) (Fig. 3h, Table S3).

### RL22’s VOC composition is distinct from the other eggplant varieties

To understand how the seven eggplant varieties relate to each other based on their VOC blend compositions, we performed a PCA. This multivariate analysis clearly showed that RL22 was different from the other six eggplant varieties (Fig. 3i). First two PCs accounted for >98% of the total variation. (Z)-3-hexen-1-ol, which was >4-fold higher in RL22 than the other eggplant varieties, contributed the most in the grouping along the first PC (Fig. 3i). Benzeneacetaldehyde, which was found in high concentrations in all seven varieties, contributed the most in the grouping along the second PC (Fig. 3i).

### RL22-specific geraniol repels ovipositing females

Since the GC analysis showed that the SFB repellent RL22 VOC blend contained four compounds that were not detected in the CEB and three compounds in significantly higher concentrations than that in CEB, we investigated whether these compounds were responsible for repellence. This was done by individually complementing these compounds on APs (Fig. 4a). 1-Hexanol-complemented APs showed >2-fold increased oviposition than the control (Fig. 4b). Complementation of (E)-3-hexen-1-ol (Fig. 4c), (Z)-3-hexen-1-ol (Fig. 4d), (Z)-3-nonen-1-ol (Fig. 4e), guaiacol (Fig. 4f) and eugenol (Fig. 4g) did not influence oviposition. Geraniol complemented APs showed no oviposition (Fig. 4h), indicating that geraniol was the repellent from RL22. Consequently, we tested whether geraniol retains its oviposition deterrent activity when it is complemented on the leaves of seven eggplant varieties (Fig. 4i); >90% reduction in oviposition was observed on geraniol-complemented plants (Fig. 4j-p). Remarkable reduction in a number of laid eggs took place in the most susceptible varieties JK (Fig. 4m) and CVK (Fig. 4p) (JK: 26.95± 2.78 in geraniol-deplete control and 1.35± 0.24 in geraniol-complemented plants; CVK: 28.4± 2.93 in geraniol-deplete control and 0.95± 0.22 in geraniol-complemented plants) (Fig. 4, S10). In all the assays, females laid more eggs on geraniol-free controls than the geraniol-complemented leaves (Fig. S4a). RL22’s geraniol concentration was found to be optimum for deterring the oviposition (Fig. S4b-c).

**Fig. 4.**
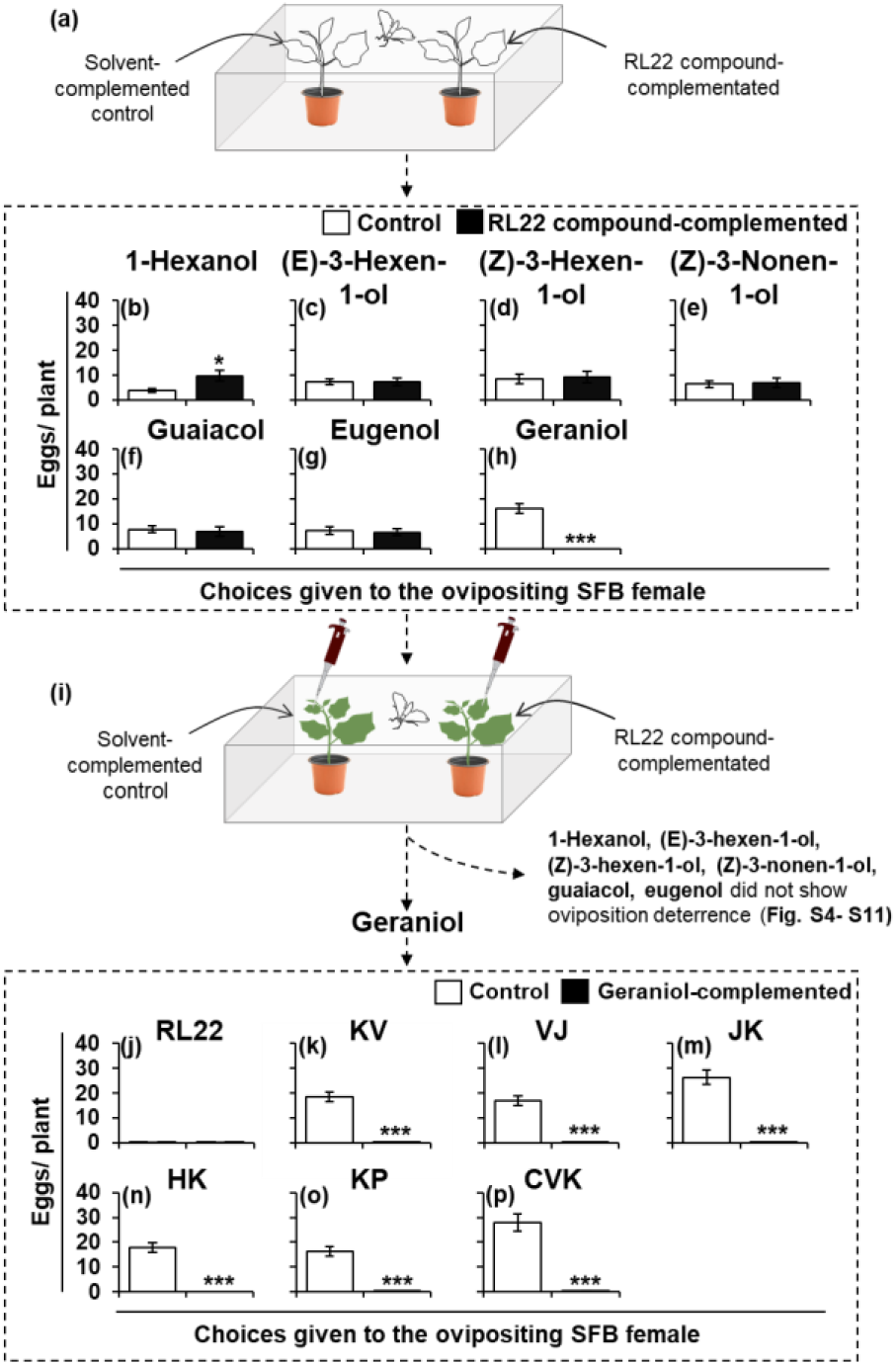
Geraniol deters oviposition. (a) A schematic of dual-choice assays conducted to analyze ovipositing SFB (*Leucinodes orbonalis*) females’ host preference using control (solvent-complemented) and RL22 compound-complemented APs. Eggs (mean±SE;) laid on (b) 1-hexanol-, (c) (E)-3-hexen-1-ol-, (d) (Z)-3-hexen-1-ol-, (e) (Z)-3-nonen-1-ol-, (f) guaiacol-, (g) eugenol- and (h) geraniol-complemented APs and their respective controls, showing that only geraniol deterred oviposition. (i) A schematic of dual-choice assays conducted to analyze ovipositing females’ host preference using control (solvent-complemented) and geraniol-complemented eggplant (*Solanum* melongena) saplings. Eggs (mean±SE;) laid on control (solvent-complemented) and geraniol-complemented (j) RL22, (k) SB, (l) NSB, (m) JK, (n) HK, (o) KP and (p) CVK, showing that geraniol complementation significantly decreases oviposition on the susceptible varieties. Asterisks indicate significant differences calculated using Student’s two-tailed t test (*≡*P*< 0.05; **≡*P*< 0.001; n= 20 assays).

1-Hexanol showed no effect on oviposition when complemented on the leaves of RL22 (Fig. S5a), KV (Fig. S5b), VJ (Fig. S5c) and JK (Fig. S5d). Similar to its complementation on AP, 1-hexanol complementation on the leaves of HK (Fig. S5e), KP (Fig. S5f) and CVK (Fig. S5g) showed >2-fold increased oviposition than the controls. Complementation of (E)-3-hexen-1-ol (Fig. S6a-g), (Z)-3-hexen-1-ol (Fig. S7a-g), (Z)-3-nonen-1-ol (Fig. S8a-g), guaiacol (Fig. S9a-g) and eugenol (Fig. S10a-g) on the leaves of seven eggplant varieties did not influence oviposition.

Solvents of these compounds water (geraniol, 1-hexanol and guaiacol) (Fig. S11a), 0.001% ethanol [(E)-3-hexen-1-ol, (Z)-3-hexen-1-ol and (Z)-3-nonen-1-ol] (Fig. S11b) and 0.1% ethanol (eugenol) (Fig. S11c) did not influence females’ oviposition.

### Silencing geraniol synthase renders RL22 susceptible to SFB oviposition

To confirm that RL22’s repellent nature is due to its geraniol content, we generated *Sm*GS-silenced RL22 plants using VIGS (Fig. 5a). These plants showed >20-fold lower *Sm*GS transcript abundance (0.05± 0.02) than that in EV (1.33± 0.27) and WT (1.03± 0.3) controls (Fig. 5b). Viral coat protein transcripts could be detected in the infiltrated *Sm*GS and EV plants, in which they showed similar levels; these transcripts could not be detected in the un-infiltrated WT plants (Fig. S12a). Transcript abundance of *Sm*MTPS1 (Fig. S12b) and *Sm*MTPS2 (Fig. S12c) did not vary in WT, EV and *Sm*GS-silenced plants, indicating that the strategy to include *Sm*GS’s unique 3’-UTR sequence in the silencing construct was successful. We inferred that the *Sm*GS silencing was sufficiently specific.

**Fig. 5.**
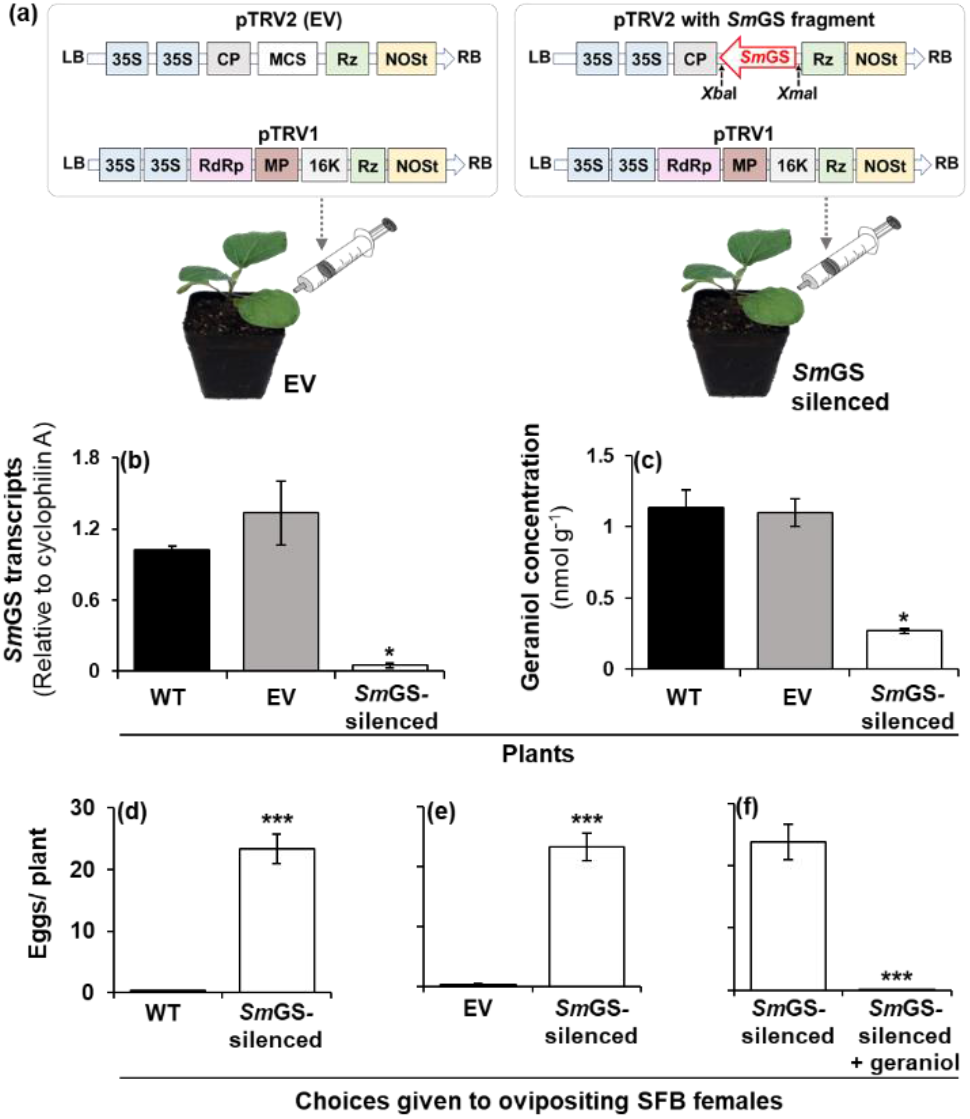
Silencing of geraniol synthase (*Sm*GS) renders RL22 susceptible to SFB (*Leucinodes orbonalis*) oviposition. (a) Schematic of constructs and infiltration procedure for virus-induced gene silencing (VIGS). (b) *Sm*GS transcript abundance (relative to cyclophilin A) in the leaves of wild type (WT), empty vector-infiltrated (EV) and *Sm*GS silencing construct-infiltrated (*Sm*GS-silenced) eggplants (*Solanum melongena*) (*F*= 369.3, *df*= 7.527, *P*< 0.0001). (c) Geraniol concentrations (mean± SE) in WT, EV and *Sm*GS-silenced RL22 leaves (*F*= 53.12, *df*= 6.959, *P*< 0.0001). Eggs (mean± SE) laid in dual-choice assays on (d) WT and *Sm*GS-silenced, (e) EV and *Sm*GS-silenced and (f) *Sm*GS-silenced and geraniol-complemented *Sm*GS-silenced RL22 plants, showing that *Sm*GS silencing renders RL22 susceptible to SFB oviposition and geraniol complementation in *Sm*GS-silenced plants restores the resistance. Asterisks in (b) and (c) indicate significant differences (*P*≤ 0.05) determined using Welch F test with Games-Howell *post hoc* test (*≡*P*< 0.05) and in (d), (e) and (f) indicate significant differences calculated using Student’s two-tailed t test (***≡*P*< 0.001; n= 20 assays).

Geraniol content of these *Sm*GS-silenced plants (14.63± 3.15 nmol g^-1^) was about 3-fold lower than that of EV (46.56± 5.75 nmol g^-1^) and WT (49.53± 8.73 nmol g^-1^) controls (Fig. 5c and Fig. S12d-f). In choice assays, SFB females preferred to oviposit on these *Sm*GS-silenced plants; ~98% of the oviposition occurred on these plants when they were used in dual-choice assays with the control WT (Fig. 5d) or EV (Fig. 5e). Lastly, the repellence could be restored by coating geraniol on *Sm*GS-silenced plants as the number of eggs laid on these plants (0.1± 0.06) was <1% of that on *Sm*GS-silenced geraniol-deplete plants (23.8± 2.78) (Fig. 5f).

## Discussion

Several studies have shown that the extent of SFB infestation varies in different eggplant varieties (Dadmal *et al*., 2004; Elanchezhyan *et al*., 2008; Javed *et al*., 2011; Devi *et al*., 2015; Kumar *et al*., 2017). Some investigators searched for the resistance factors among the physicochemical properties of the varieties showing relatively less infestation. They reported morphological characters like spines (Shaukat *et al*., 2018), trichomes (Panda & Das, 1974; Javed *et al*., 2011) and silica deposition (Kale *et al*., 1986) and contents of metabolites such as steroidal alkaloids (Doshi *et al*., 1998; Preneetha, 2002; Jat & Pareek, 2003; Thangamani, 2003), phenolics (Kale *et al*., 1986; Asathi *et al*., 2002; Prabhu *et al*., 2009), and crude fiber (Kale *et al*.,1986) as putative SFB-resistance factors. However, most of these findings were based on correlative analyses and did not empirically validate the specific resistance factors for direct use in SFB management (Shaukat *et al*., 2018; Lalita & Kashyap, 2020).

Our results suggested that ovipositing female’s host location was driven by eggplant’s foliar factors. Females showed a similar oviposition pattern on the VOC extract complemented APs as that on plants in both field and climate chamber. This ascertained that the female’s host selection was associated with foliar VOCs and not with other features such as leaf area, shape, color and surface morphology. Similar to the observations made in their native habitat and experimental field, females showed a distinct aversion to RL22 plants and RL22 VOC extract complemented APs in various choice assays. Although VOC-mediated insect resistance is known from several plants (De Moraes *et al*., 2001; Degenhardt *et al*., 2009; Staudt *et al*., 2010), RL22 VOC blend’s deterrence potential against a tenacious pest like SFB was striking, mainly because eggplant genotype with such potential is not known.

In an attempt to find the repellence factor from RL22, we found that the RL22 VOC blend was distinctly polymorphic from other eggplant varieties. It was found to be rich in C6 and C9 alcohols and aldehydes, which frequently play an important role in mounting indirect plant defense to attract herbivore’s natural enemies (Dicke, 2009; Zitzelsberger & Buchbauer, 2015). RL22 also showed a high content of some phenolics guaiacol and eugenol, which are known to confer resistance against lepidopteran herbivores in other plant species (Suckling *et al*., 1996; Molnár *et al*., 2017). However, these compounds were unable to deter SFB females. Geraniol, the only terpenoid detected from eggplants, was found to be the SFB repellant. Repellent properties of geraniol have also been reported against other lepidopteran pests like light brown apple moth (*Epiphyas postvittana* Walker; Tortricidae) (Suckling *et al*., 1996) and cabbage looper (*Trichoplusia ni* Hübner; Noctuidae) (Akhtar *et al*., 2012). Moreover, recently it was discovered that in the fall armyworm, (*Spodoptera frugiperda* J.E. Smith; Noctuidae), geraniol reduces oviposition and also impairs embryo development (Guedes *et al*., 2020). It is known that intercropping of aromatic plants like coriander *(Coriandrum sativum* L.) (Khorsheduzzaman *et al*., 1997; Satpathy & Mishra, 2011; Singh *et al*., 2016), fennel (*Foeniculum vulgare* Mill.) (Satpathy & Mishra, 2011; Singh *et al*., 2016), lemongrass (*Cymbopogon citratus* Stapf.) (Calumpang *et al*., 2013) and marigold (*Tagetes erecta* L.) (Calumpang & Ohsawa, 2015; Bhattacharyya, 2020) with eggplant reduces the SFB incidence and therefore such intercropping has now become an important component of the integrated pest management. It is striking that geraniol is a prominent component of the VOC blends of all these crops [coriander (Bandoni *et al*., 1998; Saim *et al*., 2008; Pavlić *et al*., 2015), fennel (Galambosi *et al*., 1994; Oezcan & Chalchat, 2010), lemongrass (Anonymous, 2003; Ganjewala, 2009; Ganjewala & Luthra, 2009) and marigold (Singh *et al*., 2003; Salinas-Sánchez *et al*., 2012)]. Thus, our discovery of geraniol as an antixenosis factor is in agreement with these reports and suggests that albeit unknowingly and indirectly, this compound has already been in use for the protection of eggplant crops.

Breeding for the enrichment of resistance traits has been a commonly used method to develop pest resistance, involving germplasm screening for resistance traits, classical breeding and characterization of the new phenotypes (Rotino, 1997; Elanchezhyan *et al*., 2008; Javed *et al*.,2011; Devi *et al*., 2015). However, this method is time demanding, labor-intensive and most importantly, may have unpredictable results because of the unknown physicochemical basis of the desired traits (Kos *et al*., 2009). Finding a single compound like geraniol as a basis of resistance is rare. For the management of lepidopteran pests, whose host location and selection are often performed by their adults (Renwick & Chew, 1994), disrupting of adults’ VOC-mediated host location has been suggested as a control measure (Städler, 1994; Bruce *et al*.,2005; Kos *et al*., 2009; Bruce & Pickett, 2011). Geraniol can be used for such disruption and SFB control thereof. Geraniol is a U.S. Food and Drug Administration certified ‘generally recognized as safe’ (GRAS) food additive (Sinha *et al*., 2014; Anonymous, 2020) and therefore, can be readily incorporated into the integrated pest management for a direct application to reduce the pesticide load.

Together, this work iterates the importance of insect behavioral and chemical ecology in pest management. It emphasizes that a focus on controlling ovipositing females rather than on controlling larvae by hazardous pesticide application can be a highly useful and ecofriendly strategy. Geraniol, the SFB repellent discovered by studying the SFB resistance in native Himalayan variety, can be directly incorporated in integrated pest management; geraniol overproducing eggplants or devices designed to constantly release geraniol can be thought of as durable options. Geraniol can also be used as a selection marker for developing SFB-resistant eggplant varieties.

## Acknowledgements

Authors thank the Ministry of Human Resource Development, India for the financial support and funding for the GC-MS/FID system. RG thanks Council of Scientific and Industrial Research, India for PhD fellowship, DM thanks NAMASTE+ program for scholarship and AD thanks SERB, India for N-PDF fellowship. Authors thank Indian Institute of Science Education and Research for providing the *on campus* field site, Mr. G. Pingalkar for field support and Mr. G. Pawar for help in field experiments. DMF thanks Director, ICAR Research Complex for NEH Region, Umiam for providing necessary experimental facilities.

## Author Contributions

RG, DM, MS, DMF and SP conceived and designed the experiments. All authors performed the experiments and collected the data. RG and SP conducted the statistical analyses. All authors interpreted and discussed the results. RG and SP wrote the manuscript with inputs from all the authors; SP acquired funds, administered the project and supervised the research.

## The following Supporting Information is available for this article

**Fig. S1** SFB (*Leucinodes orbonalis*) host preference in different blocks of the randomized complete block design.

**Fig. S2** SFB (*Leucinodes orbonalis*) host preference in the field positively correlates to that in multiple-choice assays.

**Fig. S3** Eggplant (*Solanum melongena*) leaf VOC blend is mainly comprised of three groups of compounds.

**Fig. S4** Geraniol deters oviposition.

**Fig. S5** 1-Hexanol complementation attracts oviposition in three eggplant varieties.

**Fig. S6** (E)-3-Hexen-1-ol does not affect oviposition.

**Fig. S7** (Z)-3-Hexen-1-ol does not affect oviposition.

**Fig. S8** (Z)-3-Nonen-1-ol does not affect oviposition.

**Fig. S9** Guaiacol does not affect oviposition.

**Fig.** S10 Eugenol does not affect oviposition.

**Fig.** S11 Solvents do not affect oviposition preference.

**Fig.** S12 *Sm*GS silencing does not co-silence highly similar *Sm*MTPS1 *Sm*MTPS2 but reduces the geraniol content.

**Table S1** Pesticides of different chemical classes against which resistance of SFB (Leucinodes orbonalis) has been reported

**Table S2** Details of primers used for the amplification of VIGS cloning fragment and qPCRs

**Table S3** VOC composition of seven eggplant (*Solanum melongena*) varieties

