## Supplemental Information for "Chemical ecology of Himalayan eggplant variety’s antixenosis: Identification of geraniol as an oviposition deterrent against the eggplant shoot and fruit borer"

**The following Supporting Information is available for this article:**

**Fig. S1** SFB (*Leucinodes orbonalis*) host preference in different blocks of the randomized complete block design.

**Fig. S2** SFB (*Leucinodes orbonalis*) host preference in the field positively correlates to that in multiple-choice assays.

**Fig. S3** Eggplant (*Solanum melongena*) leaf VOC blend is mainly comprised of three groups of compounds.

**Fig. S4** Geraniol deters oviposition.

**Fig. S5** 1-Hexanol complementation attracts oviposition in three eggplant varieties.

**Fig. S6** (E)-3-Hexen-1-ol does not affect oviposition.

**Fig. S7** (Z)-3-Hexen-1-ol does not affect oviposition.

**Fig. S8** (Z)-3-Nonen-1-ol does not affect oviposition.

**Fig. S9** Guaiacol does not affect oviposition.

**Fig. S10** Eugenol does not affect oviposition.

**Fig. S11** Solvents do not affect oviposition preference.

**Fig. S12** *Sm*GS silencing does not co-silence highly similar *Sm*MTPS1 *Sm*MTPS2 but reduces the geraniol content.

**Table S1** Pesticides of different chemical classes against which resistance of SFB (*Leucinodes orbonalis*) has been reported

**Table S2** Details of primers used for the amplification of VIGS cloning fragment and qPCRs

**Table S3** VOC composition of seven eggplant (*Solanum melongena*) varieties

**Fig. S1 SFB (*Leucinodes orbonalis*) host preference in different blocks of the randomized complete block design.** SFB infestation (%) in fruits of seven eggplant (*Solanum melongena*) varieties in (a) Block 1 ($F$= 151.4, $df$= 10.67, $P$< 0.0001), (b) Block 2 ($F$= 17.74, $df$= 10.67, $P$< 0.0001), (c) Block 3 ($F$= 51.88, $df$= 10.67, $P$< 0.0001) and (d) Block 4 ($F$= 13.65, $df$= 10.67, $p$= 0.0002) and in shoots of seven varieties in (e) Block 1 ($F$= 6.254, $df$= 10.67, $p$= 0.005), (f) Block 2 ($F$= 5.234, $df$= 10.67, $P$= 0.01), (g) Block 3 ($F$= 4.456, $df$= 10.67, $p$= 0.017) and (h) Block 4 ($F$= 5.164, $df$= 10.67, $P$= 0.01). Oviposition preference based on leaves (%) of different varieties harboring eggs in (i) Block 1 ($F$= 76.04, $df$= 11.94, $P$< 0.0001), (j) Block 2 ($F$= 48.19, $df$= 11.74, $P$< 0.0001), (k) Block 3 ($F$= 66.77, $df$= 11.96, $P$< 0.0001) and (l) Block 4 ($F$= 166.7, $df$= 11.88, $P$< 0.0001) and eggs (mean± SE) laid per plant on different varieties in (m) Block 1 ($F$= 39.85, $df$= 11.47, $P$< 0.0001), (n) Block 2 ($F$= 17.36, $df$= 11.25, $P$< 0.0001), (o) Block 3 ($F$= 48.76, $df$= 11.75, $P$< 0.0001) and (p) Block 4 ($F$= 20.28, $df$= 11.16, $P$< 0.0001). Significant differences ($P$≤ 0.05) were determined using Welch F test with Games-Howell *post hoc* test.

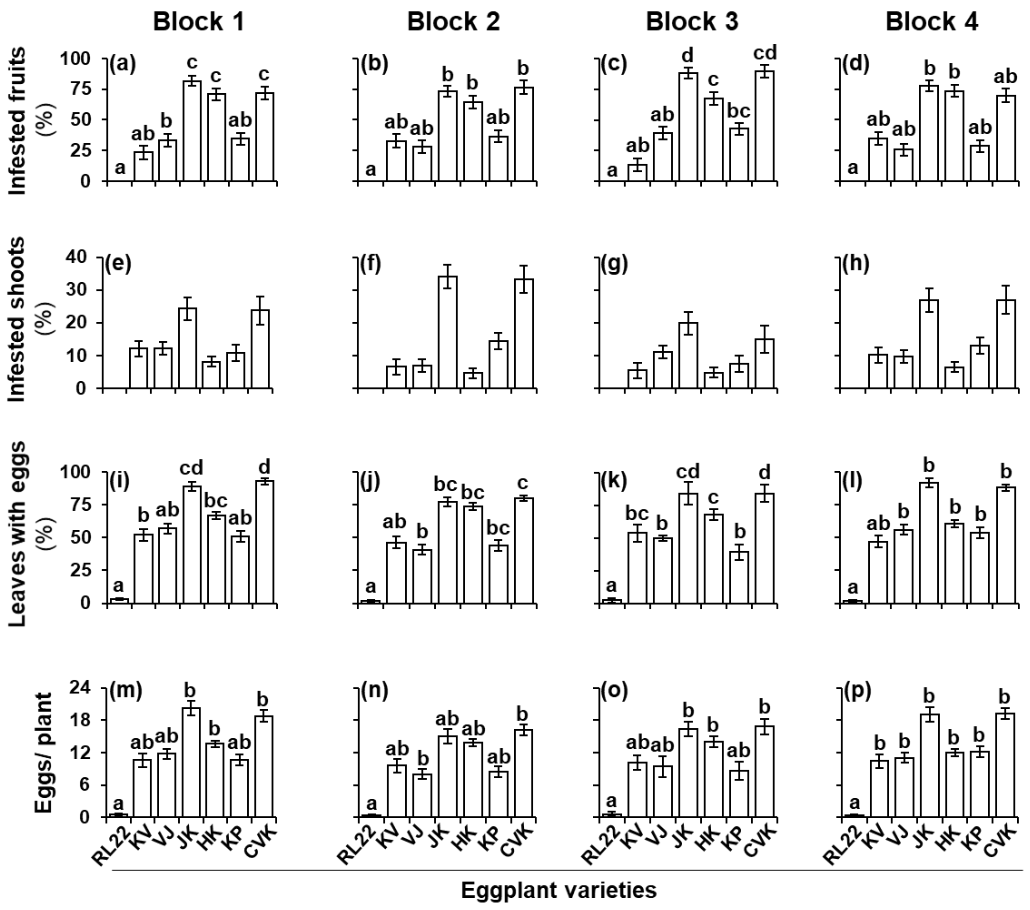

**Fig. S2 SFB (*Leucinodes orbonalis*) host preference in the field positively correlates to that in multiple-choice assays.** (a) Artificial plants (APs) made up of filter paper (grade 237 equivalent to grade 3 CHR; Sigma-Aldrich, India) used in this study. (b) Eggs (mean± SE) laid on AP and AP+ DCM in a dual-choice assay, showing that the solvent (DCM) does not affect SFB's preference. (c) Correlogram showing Spearman's ($r_s$) correlations between larval occurrence and oviposition profiles in field and oviposition profiles in the multiple-choice assays conducted using plants and VOC extract-complemented APs.

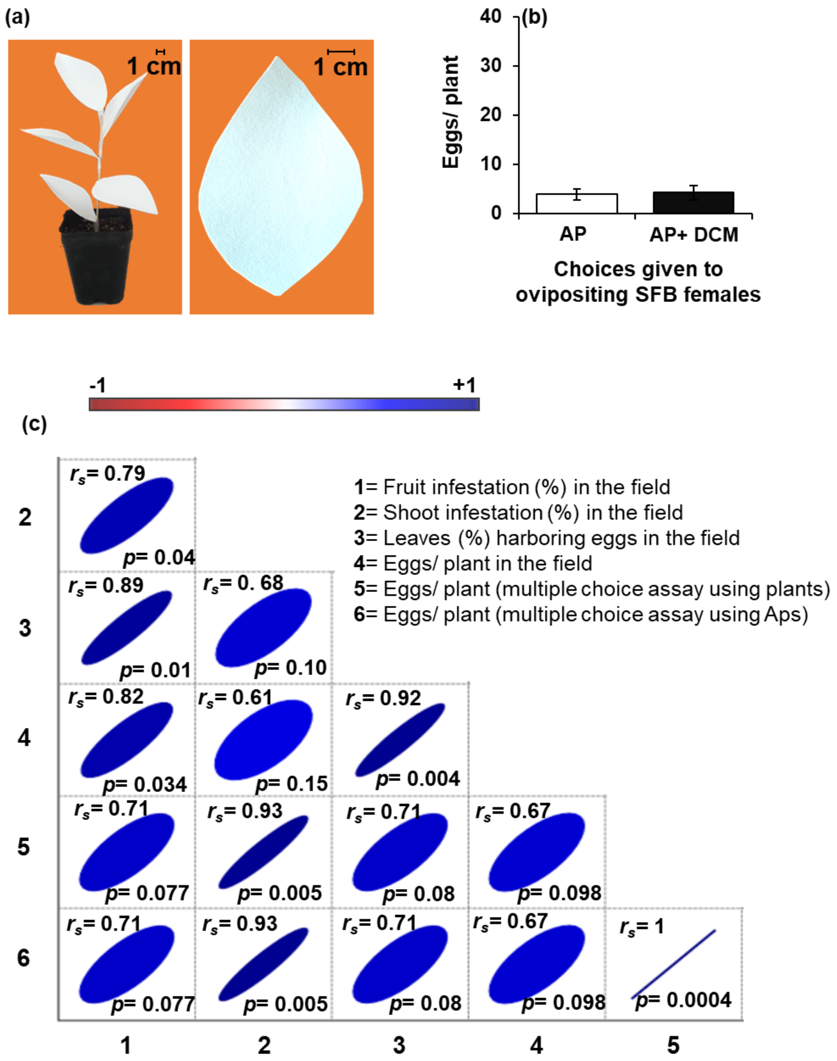

**Fig. S3 Eggplant (*Solanum melongena*) leaf VOC blend is mainly comprised of three groups of compounds.** Proportions of (a) VOCs belonging to various chemical classes, (b) benzene derivatives (c) C6 alcohols and aldehydes and (d) C9 alcohols and aldehydes in the blends of seven eggplant varieties.

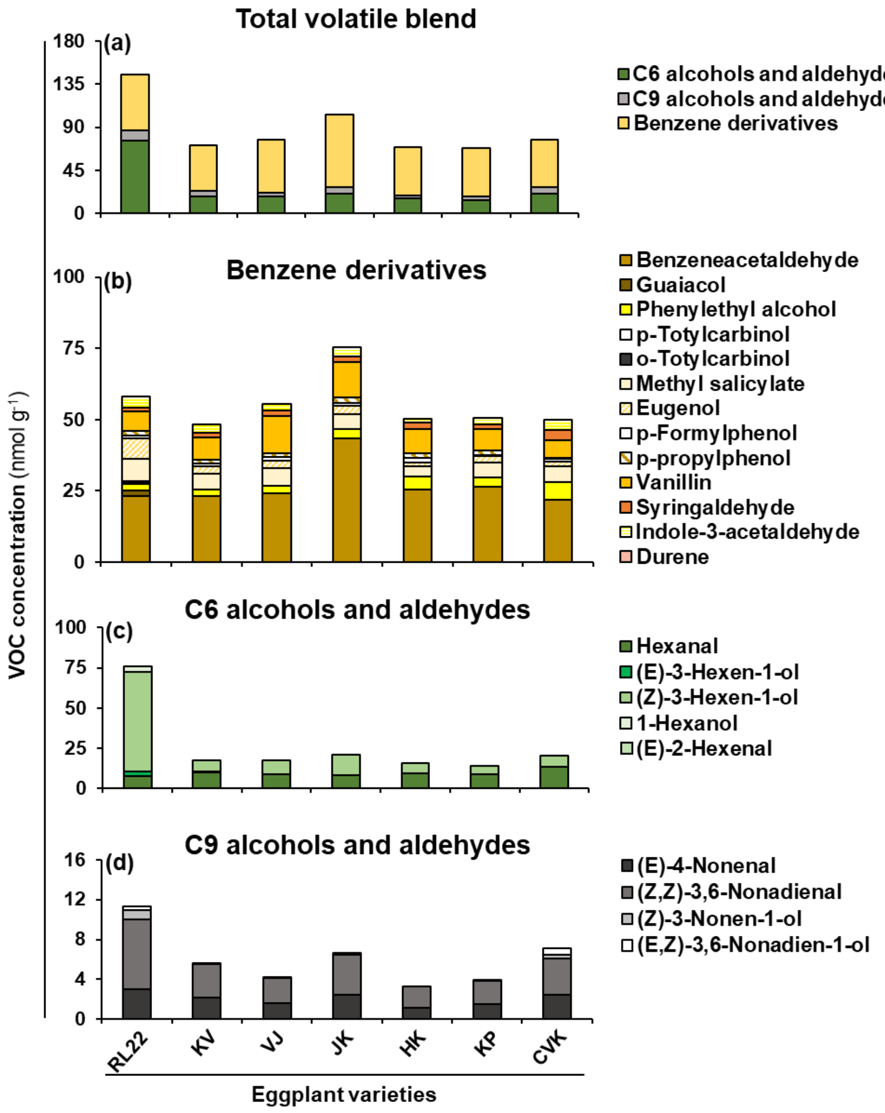

**Fig. S4 Geraniol deters oviposition.** (a) Eggs laid on control and geraniol-treated eggplant (*Solanum melongena*) leaves indicating that the geraniol complementation reduces oviposition. Eggs (mean± SE) laid on (b) CVK ($X^2 = 85.68$, $P < 0.0001$) and (c) AP ($X^2 = 65.56$, $P < 0.0001$) leaves complemented with the increasing concentrations of geraniol (uniform complementation volume of 100µl $g^{-1}$ leaf) showing that oviposition reduces with the increase in geraniol concentration (1X= 1.67 nmol $g^{-1}$, the average physiological concentration of geraniol in RL22); Significant differences ($P \leq 0.05$) were determined using Kruskal-Wallis test followed by Dunn's *post hoc* test with Bonferroni correction (n= 20 assays)

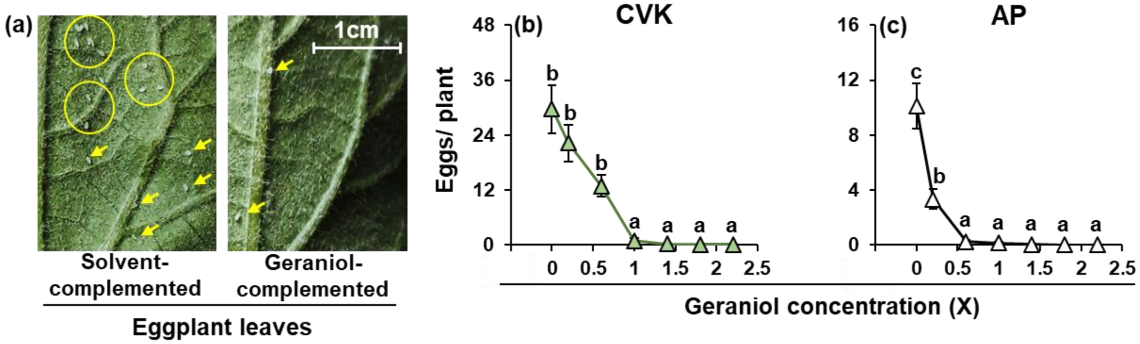

**Fig. S5 1-Hexanol complementation attracts oviposition in three eggplant varieties.** Eggs (mean± SE) laid on solvent- and 1-hexanol-complemented plants of (a) RL22, (b) SB, (c) NSB, (d) JK, (e) HK, (f) KP and (g) CVK, in dual-choice assays.

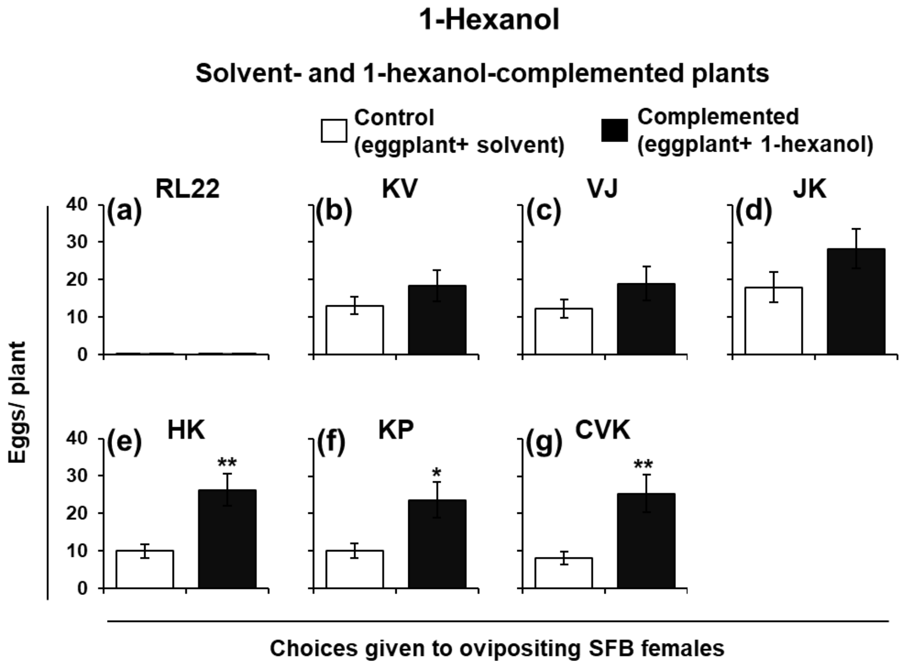

**Fig. S6 (E)-3-Hexen-1-ol does not affect oviposition.** Eggs (mean± SE) laid on solvent- and (E)-3-hexen-1-ol-complemented plants of (a) RL22, (b) SB, (c) NSB, (d) JK, (e) HK, (f) KP and (g) CVK, in dual-choice assays.

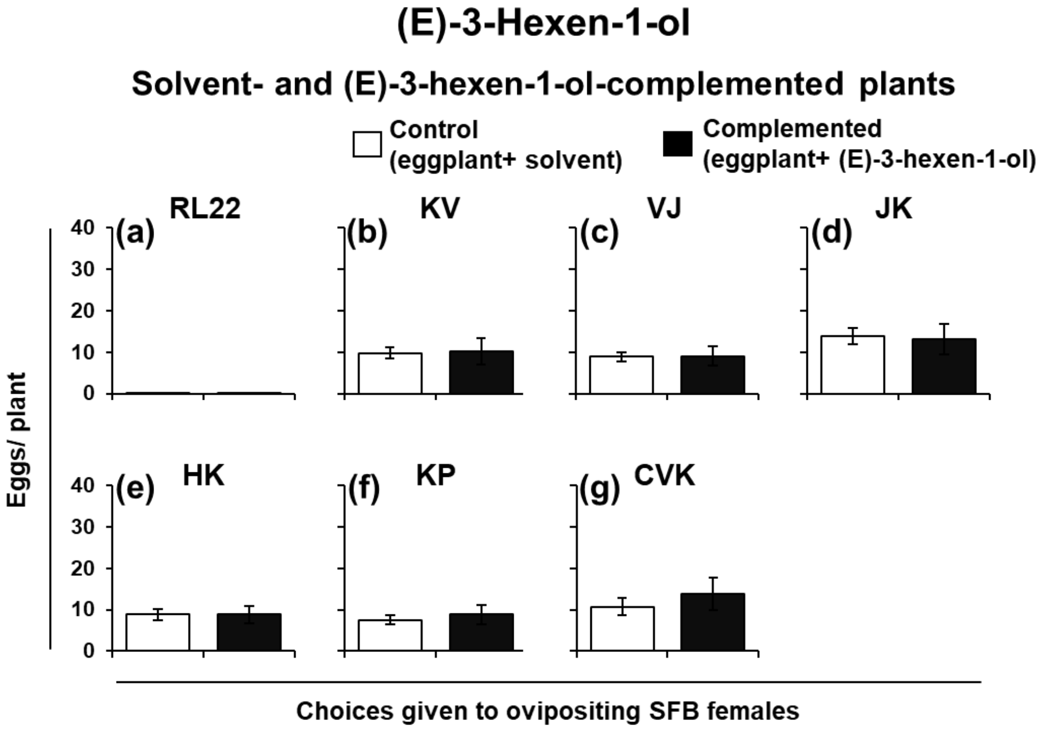

**Fig. S7 (Z)-3-Hexen-1-ol does not affect oviposition.** Eggs (mean± SE) laid on solvent- and (Z)-3-hexen-1-ol-complemented plants of (a) RL22, (b) SB, (c) NSB, (d) JK, (e) HK, (f) KP and (g) CVK, in dual-choice assays.

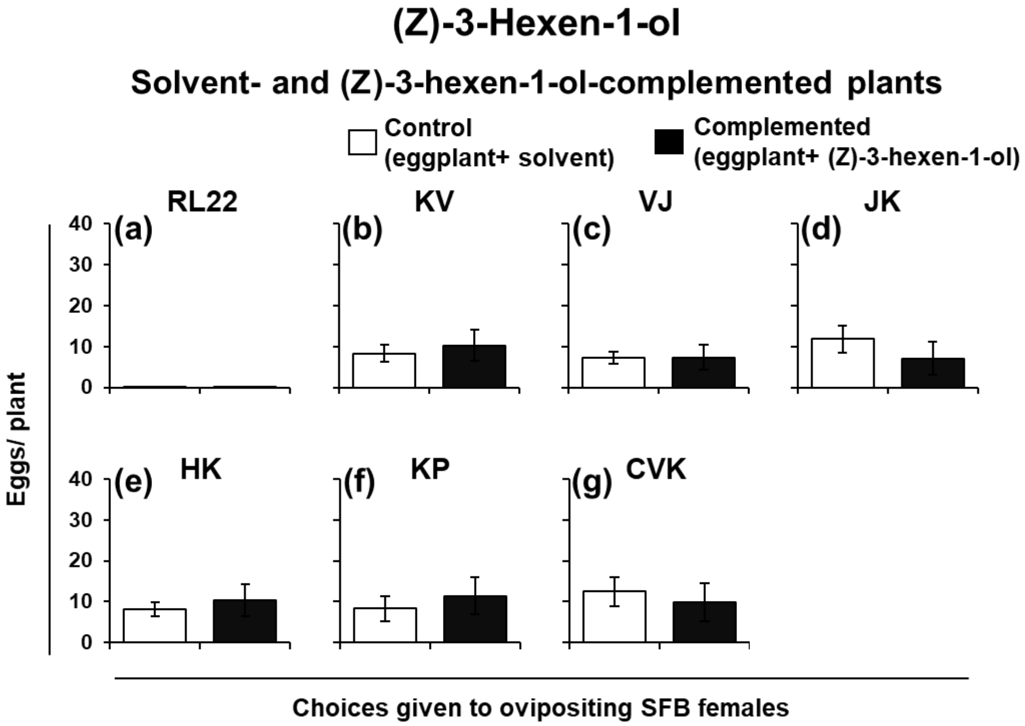

**Fig. S8 (Z)-3-Nonen-1-ol does not affect oviposition.** Eggs (mean± SE) laid on solvent- and (Z)-3-nonen-1-ol-complemented plants of (a) RL22, (b) SB, (c) NSB, (d) JK, (e) HK, (f) KP and (g) CVK, in dual-choice assays.

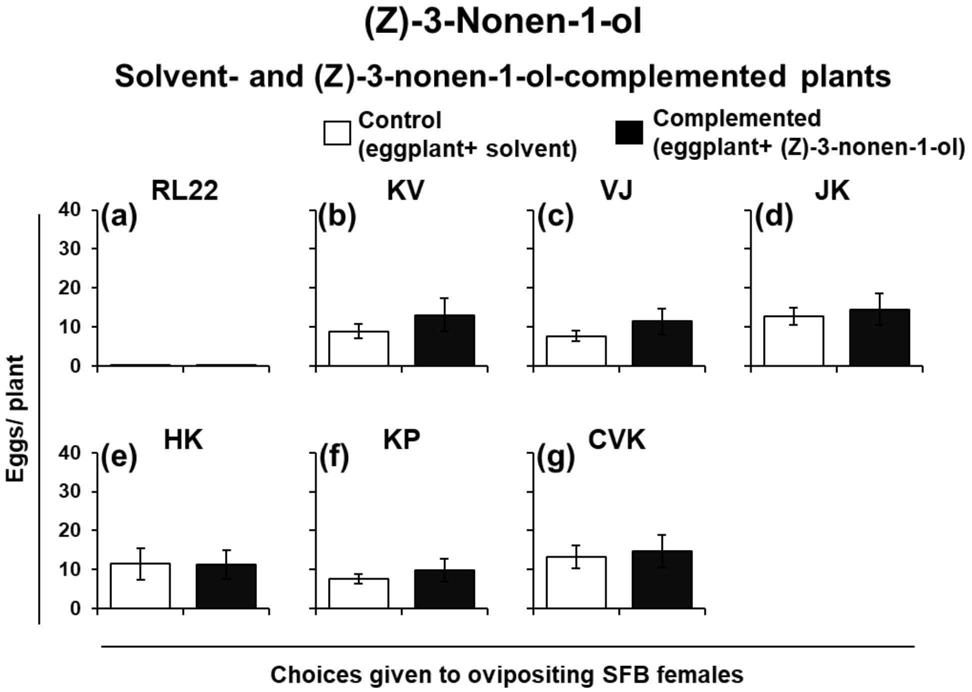

**Fig. S9 Guaiacol does not affect oviposition.** Eggs (mean± SE) laid on solvent- and guaiacol-complemented plants of (a) RL22, (b) SB, (c) NSB, (d) JK, (e) HK, (f) KP and (g) CVK, in dual-choice assays.

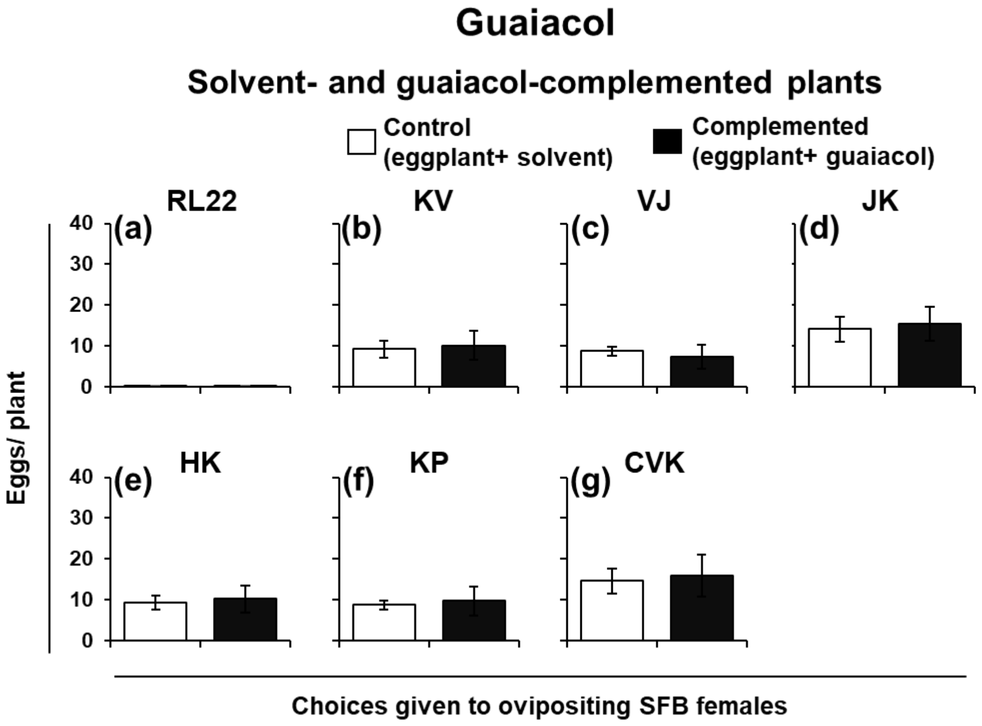

**Fig. S10 Eugenol does not affect oviposition.** Eggs (mean± SE) laid on solvent- and eugenol-complemented plants of (a) RL22, (b) SB, (c) NSB, (d) JK, (e) HK, (f) KP and (g) CVK, in dual-choice assays.

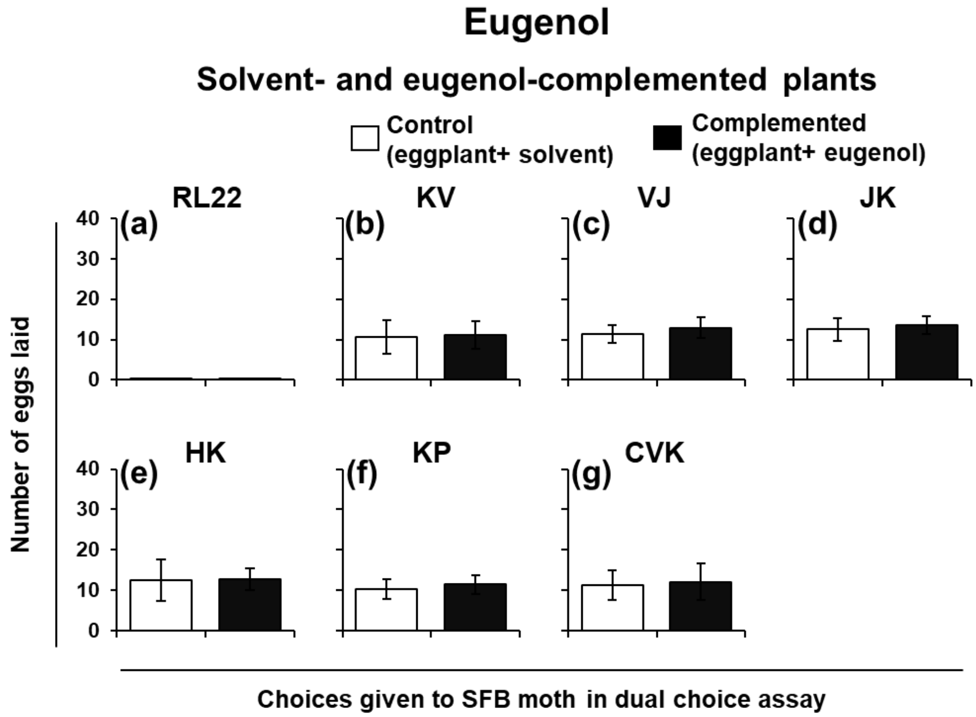

**Fig. S11 Solvents do not affect oviposition preference.** Eggs (mean± SE) laid on (a) AP and AP+ water, (b) AP and AP+ 0.001% ethanol and (c) AP and AP+ 0.1% ethanol, in dual-choice assays.

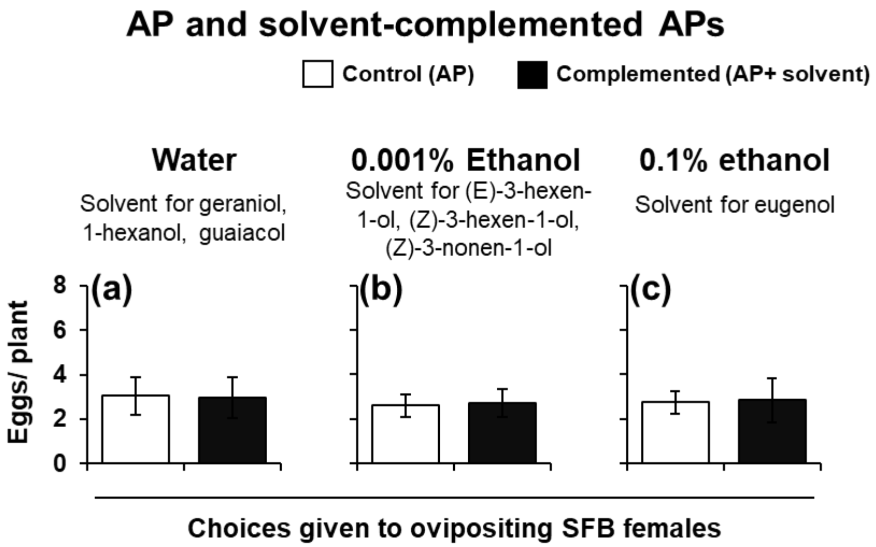

**Fig. S12 *Sm*GS silencing does not co-silence highly similar *Sm*MTPS1 *Sm*MTPS2 but reduces the geraniol content.** Transcript abundance (relative to cyclophilin A) of (a) TRV coat protein biosynthetic gene ($F$=354.8, $df$= 6.667, $P$< 0.0001), (b) *Sm*MTPS1 ($F$= 0.03, $df$= 9.954, $P$= 0.9732) and (c) *Sm*MTPS2 ($F$= 1.967, $df$= 8.87, $P$= 0.191) in the leaves of WT, EV and *Sm*GS-silenced plants. Asterisks indicate significant differences ($P \leq 0.05$) determined using Welch F test with Games-Howell *post hoc* test (*≡ $P$< 0.05). GCMS extracted ion (geraniol: m/z 69, 93,153) chromatograms of the VOC extracts of (d) WT, (e) EV and (f) *Sm*GS-silenced plants showing that the geraniol concentration remarkably reduced upon *Sm*GS silencing.

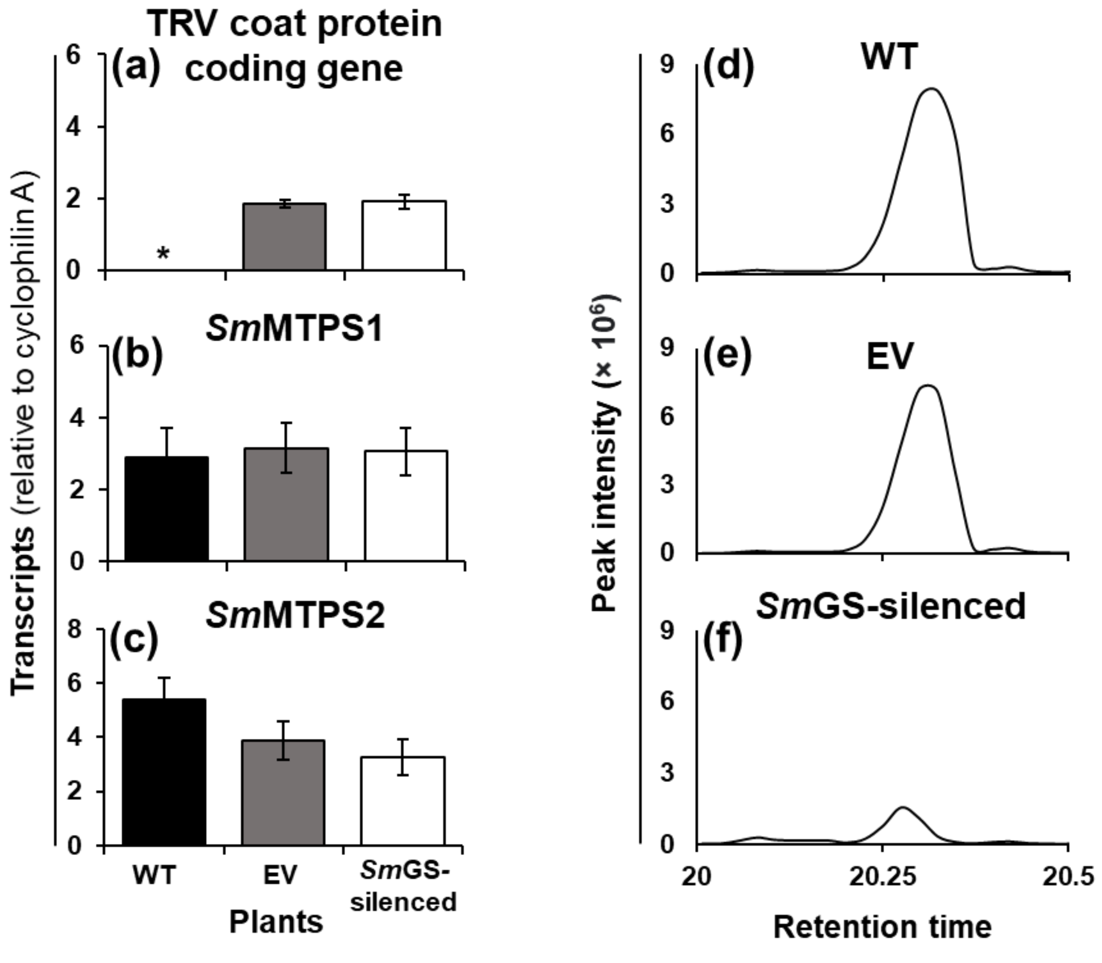

**Table S1 Pesticides of different chemical classes against which resistance of SFB (Leucinodes orbonalis) has been reported**

| **Main group and primary site of action according to IRAC (as on march 2020)** | **Sub-group or exemplifying active ingredient** | **Name of insecticides** | **Reference** |
|---|---|---|---|
| Sodium channel modulators | Synthetic pyrethroids | Cypermethrin | (Rahman, 2009) |
| | | Deltamethrin | (Murali *et al.*, 2017; Shirale *et al.*, 2017) |
| | | Alphamethrin | (Murali *et al.*, 2017) |
| | | Λ-cyhalothrin | (Murali *et al.*, 2017) |
| | | Fenpropathrin | (Murali *et al.*, 2017) |
| | | Fenvalerate | (Shirale *et al.*, 2017; Kariyanna *et al.*, 2020) |
| Gaba-gated chloride channel blockers | Organochlorines | Endosulfan | (Shirale *et al.*, 2017) |
| Acetylcholinesterase (ACHE) inhibitors | Organophosphate | Profenofos | (Shirale *et al.*, 2017) |
| | | Chlorpyriphos | (Shirale *et al.*, 2017) |
| | | Phosalone | (Kariyanna *et al.*, 2020) |
| Acetylcholinesterase (ACHE) inhibitors | Carbamate | Carbosulfan | (Rahman, 2009) |
| | | Carbaryl | (Shirale *et al.*, 2017) |
| | | Thiodicarb | (Kariyanna *et al.*, 2020) |
| Glutamate-gated Chloride channel (GLUCL) allosteric Modulators | Avermectins | Emamectin benzoate | (Botre *et al.*, 2014; Kaur *et al.*, 2014; Kariyanna *et al.*, 2020) |
| Ryanodine receptor modulators | Diamide insecticides | Cyantraniliprole | (Kodandaram *et al.*, 2017) |

| | | | |
|---|---|---|---|
| | | Chlorantraniliprole | (Botre *et al.*, 2014; Jahan *et al.*, 2018) |
| | | Flubendiamide | (Botre *et al.*, 2014; Kodandaram *et al.*, 2017; Kariyanna *et al.*, 2020) |
| Nicotinic acetylcholine receptor (NACHR) allosteric modulators-site I | Spinosyns | Spinosad | (Botre *et al.*, 2014; Kaur *et al.*, 2014) |
| Voltage-dependent Sodium channel Blockers | Oxadiazines | Indoxacarb | (Botre *et al.*, 2014) |

**Table S2 Details of primers used for the amplification of VIGS cloning fragment and qPCRs**

| Gene name | Primer orientation | Primer sequence | Used for |
|---|---|---|---|
| *Sm*GS | Forward | AAATCCCCACATAATAAGTCG | Amplification of VIGS cloning fragment |
| | Reverse | TATTGGATGAAAAGTATATCTACATCG | |
| *Sm*GS | Forward | TAGGGATGATGGTGACCTC | qPCR |
| | Reverse | GTAGGCTCGTAATTCCCTG | |
| pTRV coat protein coding gene | Forward | ACTCACGGGCTAACAGTGCT | qPCR |
| | Reverse | TCTTCCAAAGTCGAGCCAGT | |
| *Sm*MTPS1 | Forward | CGACTTCAAGGATGAGAAGGGTA | qPCR |
| | Reverse | GAGGTTTCAGAGTCCGTTGACAG | |
| *Sm*MTPS2 | Forward | TGTTCAAGCAGCACACCAACA | qPCR |
| | Reverse | CAGCCCAAAACAAATTCTCCACC | |
| Cyclophilin A | Forward | CAAAACCGCTGAGAACTTCCG | qPCR |
| | Reverse | CTTGACACATGAACCCTGGGA | |
